# HDAC1 has a role in double strand break repair by regulating γH2AX signalling

**DOI:** 10.1101/2025.11.18.689006

**Authors:** José Javier Marqueta-Gracia, Marina Bejarano-Franco, Salome Spaag, Sonia Silva, Andrés Aguilera, Belén Gómez-González

## Abstract

Genome integrity is challenged by DNA damage. DNA double-strand breaks are the most harmful DNA lesions as they block DNA replication and transcription leading to chromosome reorganisations or cell death if not properly repaired. In addition, increasing evidence points to chromatin as a relevant modulator of the efficiency of repair in eukaryotes. Here, we show that inhibition or depletion of human histone deacetylase 1 (HDAC1) regulates DSB repair by controlling the phosphorylation of H2AX, an early step of the DNA damage response. Thus, DSB repair is regulated by a crosstalk between histone acetylation and phosphorylation. Our study provides evidence that histone acetylation regulates DSB signalling supporting that histone deacetylase inhibitors could enhance genotoxic treatment of cancer.

## INTRODUCTION

The stability of the genome is challenged by DNA lesions, among which DNA double-strand breaks (DSBs) are highly cytotoxic. One of the earliest events triggered by a DSB is the phosphorylation of the histone variant H2AX at serine 139 (γH2AX), which is driven by the apical DNA damage response (DDR) kinases ATM, ATR and DNA-PK (1-4). This signal spreads to the vicinity of the DSB, serving as a platform to recruit multiple signalling and repair factors in a molecular cascade that ends in repair. DSBs can be repaired by direct re-ligation of the broken DNA ends (non-homologous end joining, NHEJ). However, DNA resection allows homologous recombination (HR), which preferentially uses the intact sister chromatid as a template (5).

To ensure faithful restoration of the genetic information, HR is restricted to post-replicative S and G2 phases of the cell cycle, when the intact sister chromatid is available (6-8). This guarantees the accurate repair of replication-born DSBs, which are a major source of DNA damage given the vulnerability of the replication process itself and the frequent stalling upon different sources of replication obstacles, including transcription and R-loops (9,10). Sister Chromatid Recombination (SCR) is favoured by the cohesion ring that holds sister chromatids together after replication (11-14) as well as by chromatin modifications and histone variants that are deposited in newly replicated chromatin, as shown in yeast for H3K56 (15,16) and in human cells for H4K20me0 (17) or the H3.3 variant (18).

Histone acetylation levels have been shown to influence the efficiency of Sister Chromatid Exchanges (SCE), which is an accurate indicator of total SCR, as shown for yeast mutations in Rpd3L and Hda1 HDACs (19). This effect was explained by a reduction in chromatid cohesion given the epistasis found between these HDAC mutations and cohesin mutants (19). Human HDACs from class I (HDAC1, 2, 3, and 8) and II (HDAC4, 5, 6, 7, 9 and 10) are related to yeast Rpd3 and Hda1, respectively (20). Among them, the class I HDAC members 1 and 2 and the scaffold factor SIN3A bind replication forks and promote their stability, thus counteracting the formation of endogenous replication-born DNA damage (21,22). Moreover, the HDAC inhibitor romidepsin was reported to cause DNA breaks through the accumulation of R-loops (22). Thus, HDAC activity is important to prevent DNA damage but, in addition, HDAC1 and 2 were shown to localize to damage sites and impact NHEJ (23). However, no major roles in HR were reported for these HDACs so far. Understanding the potential contribution of the loss of HDACs to DNA damage repair is crucial in cancer biology since histone deacetylase inhibitors (HDACi) have been shown to enhance the effect of genotoxic agents when used in combined therapy (24-27). Here we have explored the effect of human histone deacetylases in HR via the analysis of their impact on SCEs. Our results uncover a specific role for HDAC1 in SCR, not shared with the other class I HDACs, HDAC2 and HDAC3. Consequently, HDAC1 depletion impacts chromosome stability and cell proliferation upon DNA damage. We found reduced cohesin levels on chromatin, but our results rather support that HDAC1 depletion affects DSB repair by impairing the earliest step of the DSB response, namely γH2AX signalling, affecting both NHEJ and HR repair pathways. The observation that histone acetylation regulates DSB signalling, supports the view that HDAC inhibitors could be used to enhance the efficiency of cancer treatment by genotoxic agents thus exploiting the vulnerability of cancer cells, which are characterized by high levels of DNA damage.

## MATERIALS AND METHODS

### Cell cultures

Cells were grown at 37°C and 5% CO_2_ and routinely tested for mycoplasma using MycoAlert Mycoplasma Detection Kit (Lonza). For all reagents and tools, see Table S1. The U2OS cell line (ECACC, 92022711) was cultured in Dulbeccós modified Eaglés medium (DMEM) (GIBCO) supplemented with 10% heat-inactivated fetal bovine serum (FBS) (SIGMA), 2mM L-glutamine and 1% antibiotic-antimycotic (Biowest). The U2OS SEC-C cell line stably expressing a Tet-On induced Cas9 (28), was cultured under the same conditions as U2OS, but supplemented with a special FBS South America, Tetracycline Free (Biowest) and hygromycin B (100 μg/mL). The U2OS-ISce cell line, carrying a cassette with I-SceI cutting sites and a GFP reporter, was cultured under the same conditions as U2OS, but supplemented with hygromycin B (250 μg/mL). The U2OS RPA70-GFP cell line, carrying a human RPA70-GFP gene cassette, was cultured under the same conditions as U2OS, but supplemented with G418 (500 μg/mL). The BLM -/- cell line (Coriell Institute, GM08505) was cultured in Minimum Essential Medium (MEM) (GIBCO) supplemented with 10% heat-inactivated FBS (SIGMA), 2mM L-glutamine and 1% antibiotic-antimycotic (Biowest).

### siRNAs, plasmids, and gRNA transfections

siRNA (see Table S2) (50nM) transient transfections were performed twice using Lipofectamine 2000 (Invitrogen) according to manufacturer’s instructions and cells were analysed after 72 h. In U2OS SEC-C cells, 5μg/mL of doxycycline was added when indicated for 24 hours to induce Cas9-Flag expression. sgRNAs (25nM) were transfected using RNAi Max Lipofectamine (Invitrogen) according to manufacturer’s instructions and cells were analysed 6 or 24 hours later. Plasmid transfection was performed using Lipofectamine 2000 (Invitrogen) according to manufacturer’s instructions and cells were analysed after 24 hours later. Lentiviral transduction of MOI1 infective particles in U2OS-ISce cell line was performed using 10 μg/mL Polybrene (Sigma) for 24 h. Assays were performed 6 hours after adding 100nM of dexamethasone (Sigma) to induce I-SceI translocation into the nucleus.

Lentiviral pSIN-mCherry-I-SceI-GR vector for stable expression of I-*Sce*I-mCherry was generated by a cloning a PCR fragment containing the mCherry-I-SceI-GR fragment into BamHI/NotI digested pSIN-DUAL-GFP.

### Metaphase spreads for SCE, PSCS and breaks and gaps analysis

Metaphase spreads were performed as described in (29) with minor modifications. After 48 hours of the first siRNA transfection, cells were incubated with 10 μM BrdU to be incorporated into DNA for 66 and 48h respectively to allow two rounds of replication. This step was not required for the analysis of premature sister chromatid separation. When indicated, cells were irradiated with 2Gy of IR 8 hours before collection and at the same time 50mM caffeine was added to the medium to allow G2 cell cycle progression. Treatment with 0.1 μg/mL of KaryoMAX colcemid solution (3h 30min, 37°C) was then used to enrich the mitotic population before collection. Cells were detached with accutase, washed with PBS 1X, incubated in 75 mM KCl (10 min, 37°C) and followed by 5 consecutive washes with of 3:1 methanol:acetic acid. Cells were dropped onto slides covered by Acetic acid 45% and dried (o/n min, 65°C). To differentially stain the two chromatids, slides were incubated (20 min) with 10 μg/mL Hoechst 33258 (AnaSpec) in distilled water, rinsed and exposed to UVA irradiation (1 h) on a tray filled with SSC 2X (30 mM sodium citrate pH 7.0, 300 mM NaCl). Then, slides were incubated again (20 min, 60°C) in SSC 2X, rinsed and incubated in methanol (5 min, RT). Finally, cells were stained (20 min) with 5% Giemsa in Weise Buffer pH 6.8 (Sigma Aldrich), rinsed twice with Weise Buffer and allowed to dry before imaging. Data acquisition was performed with Nikon NI – SSR microscope with NIS Elements 4.0 software using a 100x objective for metaphase spread assays. Measurements were scored manually with ImageJ 1.53a software in previously blinded images.

### Western blotting

Whole-cell extracts were prepared by incubating 5 min at 95°C in Laemmli buffer 1X (50 mM Tris-HCl, 10% glycerol, 2% SDS, 0.1% Bromophenol Blue, 100 mM β-mercaptoethanol). Proteins were separated in in 4-20% gradient SDS-PAGE CriterionTM TGX TM Precast Gels (BioRad) and standard methods were used for SDS-PAGE and protein immunoblots. Ponceau S was used to check protein loading. Blocking of nitrocellulose membranes (Merk) was performed with Blocking reagent (Roche). Primary antibodies were incubated o/n at 4°C in blocking reagent and secondary antibodies 1 hour at RT. SuperSignal West Pico Plus (ThermoFisher) was used for chemiluminescence detection by ChemiDoc XRS system with ImageLab 6.0.1 software.

Protein levels from Western blot signals were quantified using the Analyze Gel tool in ImageJ software version 1.51a. The bands were selected, plotted, and the area under the curve was measured. Protein signal intensity was assessed relative to the loading control and normalized to the control sample.

### Survival assays

U2OS were seeded, and siRNA transfected. 48 hours after seeding, cells were collected trypsin (Sigma-Aldrich), scored, and seeded at same confluency (60-70%). 24 hours later, cells were exposed to 2Gy IR and 24 hours after IR, cells were collected using accutase (Sigma-Aldrich) and automatically counted by CellDrop FL (DeNovix). For each depletion, the percentage of surviving cells was calculated relative to the non-irradiated condition.

### Cell proliferation analysis by EdU incorporation

Click-iT EdU Imaging Kit (Invitrogen) was used to assay the newly synthetized DNA *in vivo* during replication. 30 min before IR, U2OS cells were incubated with EdU (10 μM). Time 0 cells (15 min after IR) coverslips were collected, and time 8 (8 hours after IR) cells were maintained with EdU to allow DNA damage recovery. Samples were fixed, permeabilized and Click-iT reaction (Alexa Fluor azide 647) performed according to the manufacturer’s guidelines. Coverslips were then washed twice and blocked with PBS-BSA 3% for 1 h, RT and darkness and continued with the immunofluorescence.

### Immunofluorescence

Cells were collected using trypsin (Sigma-Aldrich) and fixed in 4% formaldehyde in PBS 1X for 10 min at RT. Then cells were washed three times with PBS 1X and permeabilized with 70% ethanol for 5 min at -20°C followed by 5 min at 4°C. Cells were washed again three times in PBS 1X prior blocking step. Cells were blocking using with PBS-BSA 3%. Afterwards, the coverslips were incubated with primary antibodies diluted in PBS-BSA 3%. Then, cells were washed three times in PBS 1X and incubated with secondary antibodies conjugated with Alexa Fluor in PBS-BSA 3% for 1 hour at RT in darkness. Next, cells were washed twice with 1X PBS and nuclei were counterstained with 1 μg/ml DAPI in PBS 1X for 5 min in darkness, washed three times in 1X PBS, once in distilled water and mounted with a drop of ProLong Gold antifade reagent (ThermoFisher) before imaging.

Data acquisition was performed with LAS AX (Leica) microscope equipped with a DFC390 camera and 63x objective. Measurements were acquired and processed by automated scoring using MetaMorph v7.5.1.0. software to avoid bias.

### Alkaline comet assay

Comet assay was performed using a commercial kit (Trevigen) following the manufacturer’s protocol. Cells were collected using accutase, counted and resuspended at 200,000 cells/mL per ml in ice cold PBS 1X. Then cells were mixed with pre-warmed (37°C) low melting agarose and carefully dropped onto on CometSlides in flat (30 min, 4°C). From this point, all steps were performed in darkness conditions. Then slides were immersed in the commercial lysis solution from the kit (30 min, 4°C) and buffer excess was drained. Then, DNA was unwound and denatured in freshly prepared alkaline unwinding solution pH>13 for 20 min at RT. Next electrophoresis was performed in prechilled (4°C) alkaline electrophoresis for 30 min. After electrophoresis solution excess was drained, slides were immersed twice in distilled H_2_O for 5 min each, then in 70% ethanol for 5 min and dried at RT. DNA was stained with SYBR Gold Nucleic Acid Gel Stain 1X (Invitrogen) at 4°C for 5 min and slides were allowed to dry before imaging.

Data acquisition was performed with LAS AX (Leica) microscope equipped with a DFC390 camera and a 10x objective. Measurements were acquired and processed by TriTek CometScore Professional v 1.0.1.60 software. Tail moment reflects both the tail length and the fraction of DNA in the comet tail (Tail moment = (DNA in tail * tail length) / 100).

### Laser microirradiation-induced DNA damage

24 hours before microirradiation, cells were collected and seeded in μ-slide 8 well glass bottom (Ibidi) for *in vivo* cell imaging and μ-slide 8 well Grid 500 (Ibidi) for immunofluorescence after microirradiation. When necessary, plasmid transfection with the reporter proteins were performed in parallel with the seeding. DMEM medium was exchanged for pre-warmed Leivobitz media (GIBCO) just before microirradiation. Microirradiation took place in a spinning disk confocal microscope (Zeiss) associated with a computer and control systems (GATACA systems) previously equilibrated to 37°C and 5% CO_2_ with the 100x objective. For *in vivo* cell imaging, one laser stripe inside the nucleus was performed. A pre-laser image was taken prior microirradiation and sequential images along the time after the microirradiation allowed to see protein fused to GFP recruitment at damaged site. Immunofluorescences assays after microirradiation were performed as in (30) with some minor modifications. A set of horizontal and parallel laser stripes in different fields were performed. Then, media was removed, and cells washed carefully once with cold PBS 1X. Pre-permeabilization buffer was added for exactly 2 minutes and in the fume hood, the buffer was removed and, without washing cells, fixative solution was added for 15 minutes at RT in oscillation to ensure proper mixing of the fixation buffer. Cells were wash three times with PBS 1X during 10 min each wash and treated with the permeabilization buffer 10 minutes at RT in oscillation. Cells were washed again twice with PBS 1X prior blocking step of immunofluorescence.

Data acquisition for laser microirradiation immunofluorescence was performed with Thunder DMi8 (Leica) microscope equipped with a KI5 camera, using a 63x objective. The sequential images obtained from live-imaging or after immunofluorescence were analysed by ImageJ2 v2.9.0/1.53t software considering three defined regions of interest (ROIs) with the same area: i) laser ROI: Laser incision regions, ii) cell ROI: located inside the cell nucleus, except in the laser incision region, and iii) background ROI: located outside the cell in an area without fluorescence as background noise. Mean gray value is obtained with FIJI and the value for each timepoint or cell is calculated with the following equation:

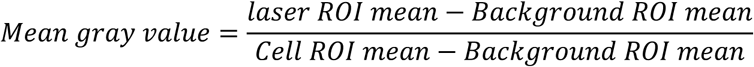

### Chromatin immunoprecipitation (ChIP) and qPCR analysis

ChIP experiments were performed according to (31) with minor modifications. Briefly, cells were crosslinked and resuspended in cell lysis buffer 1, followed by centrifugation. The resulting pellet was then treated with nuclei lysis buffer 2. Chromatin was sonicated in 2 cycles of eight pulses 30” ON/OFF in Bioruptor (Diagenode), to obtain <1kbp fragments. For each sample, 30 μg of chromatin was diluted 1/10 with IP buffer. Chromatin were incubated o/n at 4°C with ChIP-grade antibodies. A negative control with the same amount of IgG from rabbit or mouse antibody was used to calculate the background signal. Chromatin-antibody complexes were immunoprecipitated for 2 hours with 30 μl of Dynabeads Protein A (Invitrogen) at 4 °C and washed once with wash buffer 1, once with wash buffer 2, once with wash buffer 3 and twice with 1X TE. Input and immunoprecipitated samples were then de-crosslinked in elution buffer at 65 °C o/n. Finally, after phenol-chloroform purification, DNA was resuspended in 50 μL of milliQ H_2_O.

Real-time quantitative PCRs (qPCRs) were performed on a 7500 Fast Real-Time PCR system (Applied Biosystems). Results were analysed with 7500 System Software V2.0.6. Oligonucleotide used are listed in Table S2. The percentage of input recovered for each immunoprecipitated sample was calculated using the following equation: 100 × 2^*CT*^ ^*IP*-[*CT*^ ^*input*-(*log*2^ ^*DF*)]^ where CT is cycle threshold and DF is dilution factor (1:10). For relative amount quantification, The relative fold change respect to control condition was calculated using the following ΔΔ*CT* equation: 2^&[(Δ*CT*^ ^*sample*-Δ*CTcontrol*)^ where Δ*CT sample* is the difference between CT values of each sample from the region of interest (near the DSB) and region control (outside the DSB) and Δ*CT control* is the Δ*CT sample* for siC condition.

### Quantification and statistical analysis

Statistical significance analysis was performed using GraphPad Prism software v10.0.3. The number of biological replicates, cells analysed, statistical tests used, and corresponding p-values are provided in the figure legends. Sample selection and imaging fields for immunofluorescence were chosen at random. Unless stated otherwise, data collection and analysis were carried out automatically using dedicated software to minimize bias. Biological replicate variation is expected to be normally distributed and exhibit equal variance. Student’s t-test, Mann-Whitney test and Two-way ANOVA test were used when data meet the assumptions of the selected test. P values <0.05 were considered as statistically significant (****, p< 0.0001; ***, p< 0.001; **p, <0.01; *p <0.05).

## RESULTS

### HDAC1 is required for SCEs in BLM-/- cells independently of sister-chromatid separation

To determine whether human HDACs have a role in SCR, we tested the effect of class I and class II HDAC inhibitors in BLM-/- cells, which are characterized by high levels of SCEs due to impaired dissolution of double Holliday junctions resulting from the repair of spontaneous DNA lesions (32-34). Trichostatin A (TSA) was used as a pan-inhibitor of class I and II HDACs, and romidepsin as a more specific inhibitor, targeting HDAC1, HDAC2 and partially HDAC3 (35). We observed an average of 1.15 SCEs per chromosome per metaphase in control conditions (DMSO) that was reduced to 1 after treatments with either TSA or romidepsin (Figure 1). Since this result points to a role of HDACs targeted by romidepsin in SCE, we assessed the effect of depleting either HDAC1, 2 or 3 with siRNAs. Interestingly, we found that depletion of HDAC1, but not HDAC2 or HDAC3, caused a significant reduction in SCE levels, suggesting that HDAC1 might affect the repair of DSBs by HR. As a control, we confirmed that depletion of the SCC1 cohesin component decreased SCEs, in agreement with the reported role of cohesins in SCR (13,14).

**Figure 1.**
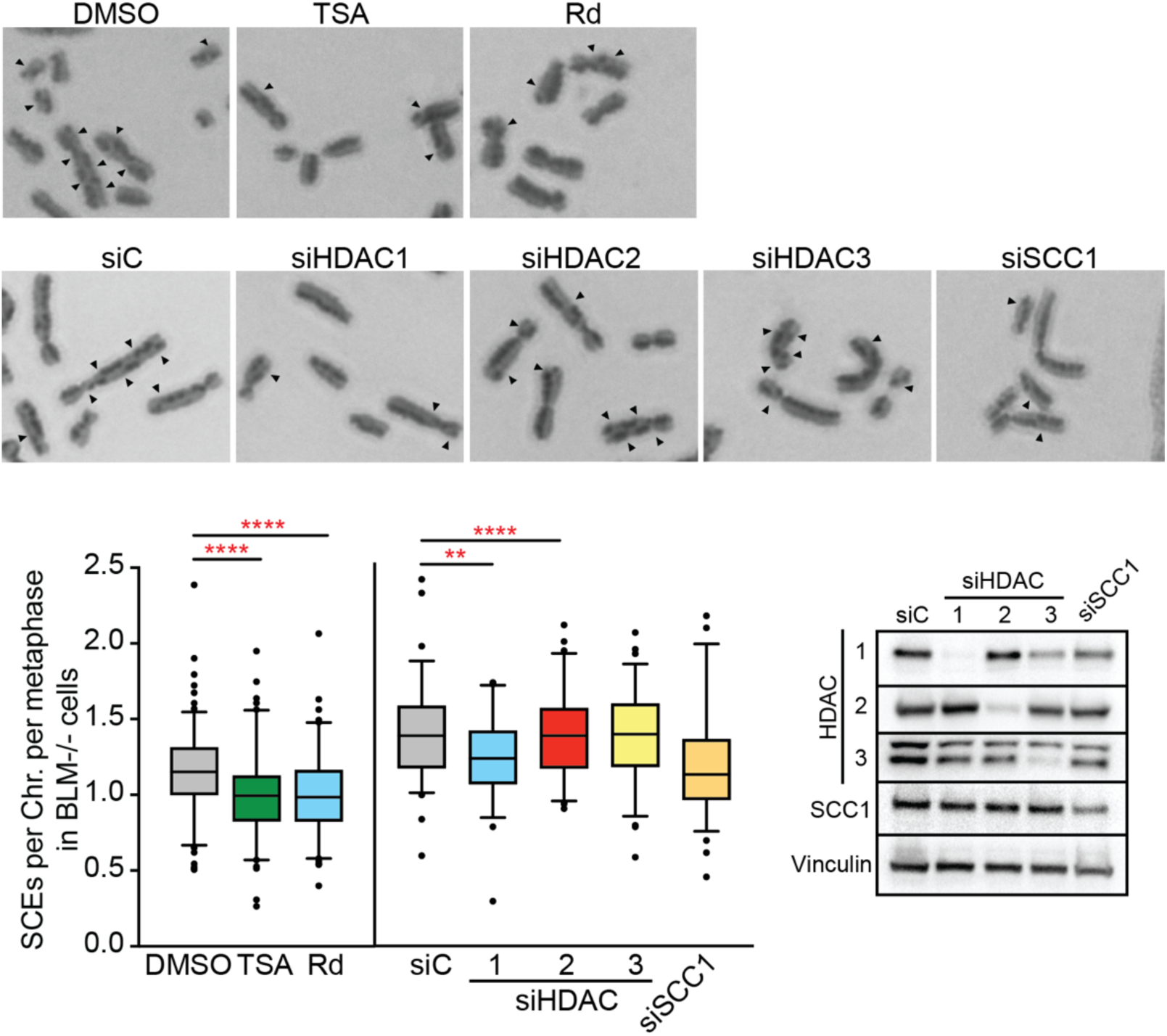
Histone deacetylation is required for SCE in BLM-/- cells. Images of HDACi-treated or siRNA- depleted BLM-/- metaphase spreads with differentially stained chromatids. Arrows indicate exchanges between chromatids. Box and whiskers (5-95 percentile) plots show distribution of SCEs per chromosome and metaphase. 25 metaphases analysed per experiment. Data pooled from 3 experiments. ****, p<0.0001; two- tailed Mann-Whitney test. Representative Western-Blot with anti-HDAC1/2/3, anti-SCC1 and anti-vinculin antibodies.

To explore the mechanisms by which HDAC1 could be affecting SCR, we first determined the impact of depleting HDAC1 on sister chromatid cohesion. Cohesin levels were measured by SCC1 chromatin immunoprecipitation (ChIP) at three different previously reported cohesin binding sites, CBS2, 3 and 9, which locate at chromosomes 5, 6 and 22, respectively (36,37). As expected, cohesin depletion by siSCC1 significantly reduced ChIP signals at all cohesin binding sites tested (Figure S1A). HDAC1 depletion also decreased cohesin levels at the three CBSs analysed, but the decrease was only significant for CBS3, consistent with a mild cohesion defect. To evaluate the effect of HDAC1 depletion in global cohesion levels, we measured premature sister-chromatid separation, known to increase upon major cohesin loss (38). SCC1 depletion significantly increased the percentage of cells with a premature sister chromatid separation phenotype, as expected (Figure S1B). However, HDAC1 depletion did not cause any significant variation. Therefore, HDAC1 loss slightly impairs the loading of cohesin at CBS but not to levels sufficient to induce detectable premature sister-chromatid separation in mitosis. We therefore conclude that any possible effect of siHDAC1 on cohesion loading does not explain by itself its effect in DSB repair by SCR.

### HDAC1 is required for SCE and cell survival upon IR

To test whether the effect of HDAC1 inhibition or depletion on SCEs is general and not specific for BLM-/- cells, we analysed SCE in U2OS cells after Ionizing Radiation (IR) (Figure 2). The spontaneous average levels of SCEs in U2OS cells were below 0.15 per chromosome per metaphase, significantly lower than that of BLM-/- cells as expected. HDAC1 depletion (Figure 2B) significantly increased SCE levels consistent with an increased incidence of spontaneous DNA damage (39,40) but more importantly, IR-induced SCEs were significantly reduced by both romidepsin and HDAC1 depletion (Figure 2A and B). This supports that HDAC1 loss affects repair of DSBs by HR also in U2OS cells and regardless of whether SCEs are induced via BLM inactivation or IR. Notably, and in agreement with defective DNA break repair, we observed significantly more breaks and gaps upon IR in HDAC1-depleted cells than in the U2OS control (Figure 2C). Consequently, we assessed whether this HR defect affected cell survival upon IR, as would be expected. The percentage of survival 24 hours after IR was significantly reduced to half in HDAC1-depleted cells, but not in wild-type control cells (Figure 2D). Altogether, these data support that HDAC1 is required for DSB repair by HR, and consistently, HDAC1 depletion leads to unrepaired DNA breaks and gaps and loss of viability upon IR.

**Figure 2.**
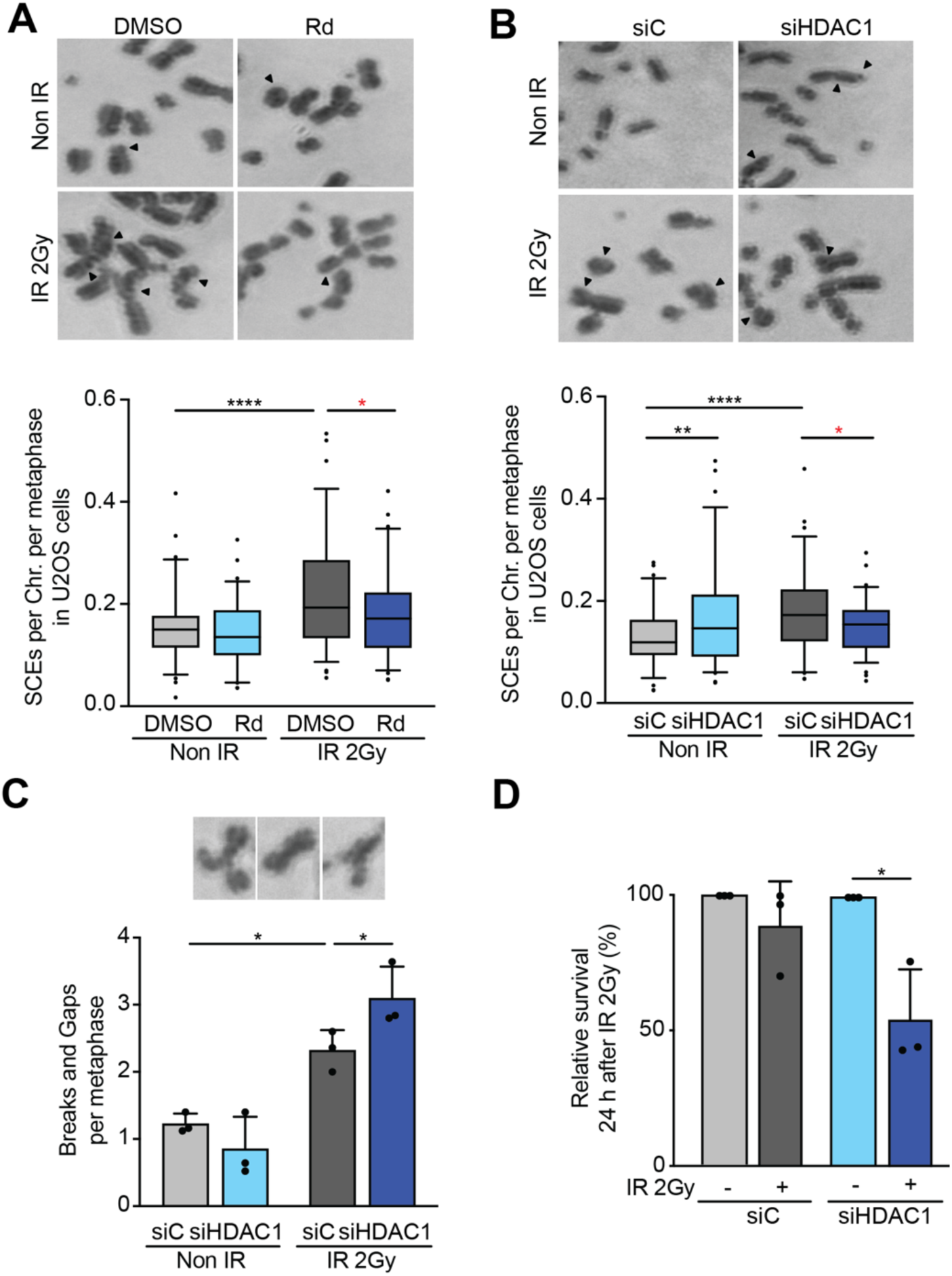
HDAC1 depletion negatively affects SCE, chromosome stability and cell survival upon IR. **(A)** Images of HDACi-treated U2OS metaphase spreads with differentially stained chromatids with or without IR. Arrows indicate exchanges between chromatids. Box and whiskers (5-95 percentile) plots show distribution of SCEs per chromosome and metaphase. 25 metaphases analysed per experiment. Data pooled from 3 experiments. ****, p<0.0001; *, p<0.05; two-tailed Mann-Whitney test. **(B)** Images of siRNA-depleted U2OS metaphase spreads with differentially stained chromatids with or without IR. Arrows indicate exchanges between chromatids. Box and whiskers (5-95 percentile) plots show distribution of SCEs per chromosome and metaphase. 25 metaphases analysed per experiment. Data pooled from 3 experiments. ****, p<0.0001; **, p<0.01 *, p<0.05; two-tailed Mann-Whitney test. **(C)** Images represent chromosome breaks and gaps of U2OS cells after IR. Histograms show the quantification of breaks and gaps per metaphase (mean±SD, N=3). siRNA as indicated. *, p<0,05; two-tailed Paired t-test. **(D)** Histograms show the percentage of relative survival (mean±SD, N=3) of irradiated U2OS cells versus non-irradiated conditions of each siRNA 24 hours after IR. *, p<0,05; two-tailed Paired t-test.

### HDAC1 is required for efficient repair of IR-induced DNA damage in G2

Cells defective in HR are known to accumulate DSBs in G2 since HR is the most effective DSB repair pathway in this cell cycle phase. The accumulation of γH2AX foci in G2 cells at late times after IR is a proxy for defective HR as it represents the percentage of unrepaired DSBs (18). Consequently, we examined γH2AX foci in U2OS cells depleted of HDAC1, 2 or 3 just after IR and 8 h later (Figure 3A and B). Quantifications were referred to G1, S and G2 phases, as determined by the DNA content (DAPI staining) and the replicating stage of each cell, measured as incorporated EdU, which was added to the medium 30 min before IR (Figure 3B and C). To estimate the repair efficiency of IR-induced DNA breaks, we calculated the ratio of γH2AX foci remaining after 8h versus those at time 0 at each stage of the cell cycle. This ratio was significantly higher in HDAC1-depleted cells versus the control only in the case of G2 cells, whereas this was not the case for siHDAC2 and siHDAC3-treated cells (Figure 3D,E and F). As a positive control, siSCC1 depletion was also analysed, leading to an increased ratio of unrepaired IR-induced DNA breaks specifically in G2 cells, as reported (41). These results support a specific role of HDAC1 in DSB repair by HR in agreement with the reduced number of damage-induced SCEs upon HDAC1 depletion or inhibition.

**Figure 3.**
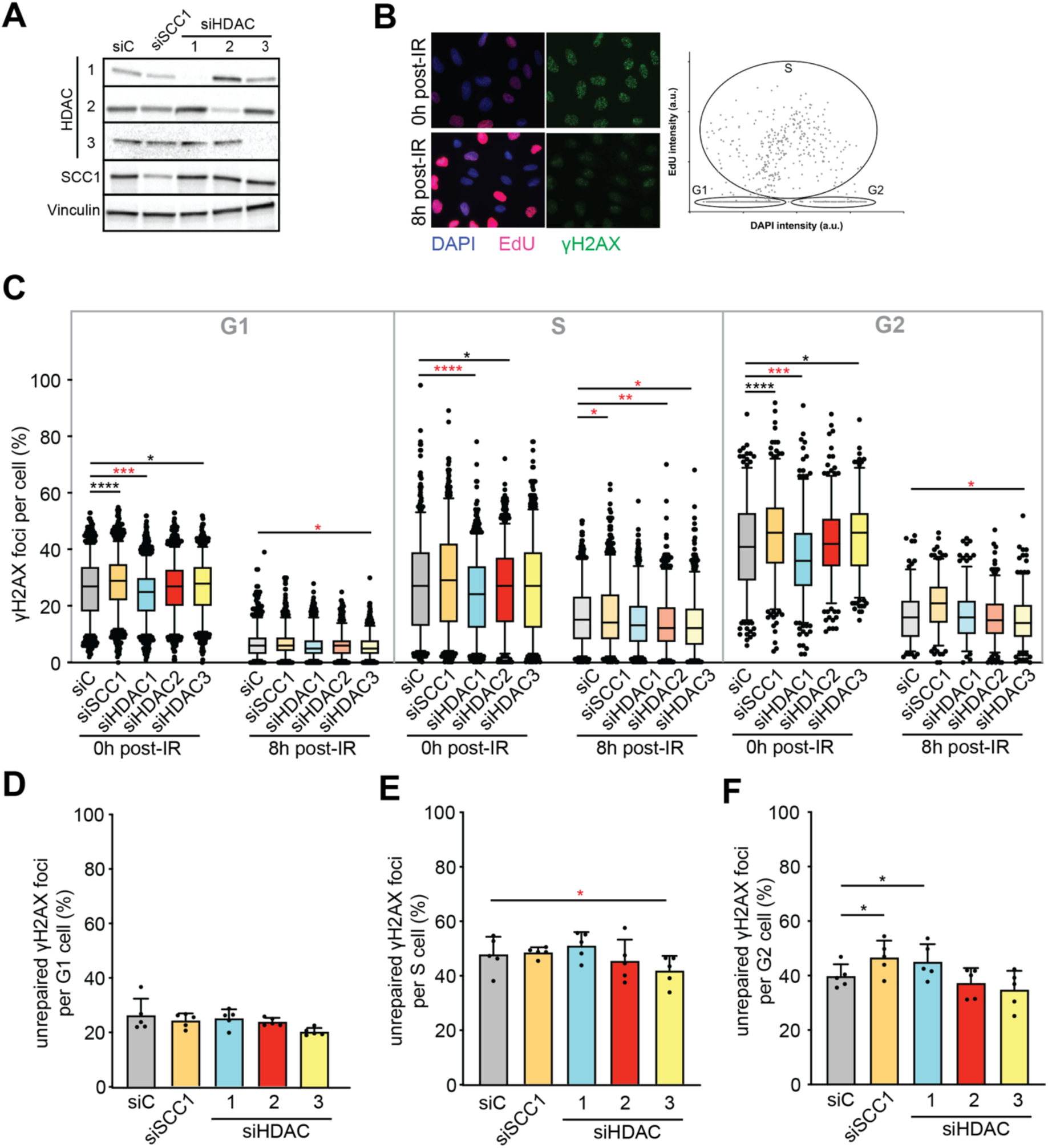
HDAC1 depletion accumulates unrepaired IR-induced DSBs in G2. **(A)** Representative Western-Blot of irradiated U2OS cells at time 0 with anti HDAC1/2/3, SCC1 and vinculin antibodies. **(B)** Immunostaining with γH2AX (green) antibody, EdU (red) and DAPI (blue) in images of siC cells. Times post IR as indicated. Scatter plot shows representative distribution of siC cells according to their DAPI and EdU content at time 0. Circles indicate the cell cycle phases. **(C)** Box and whiskers (5-95 percentile) plots show distribution of γH2AX foci per cell at G2-phase. Time 0 and 8 post IR. siRNA conditions as indicated. Data pooled from 5 experiments. 60 and 50 cells scored per condition and experiment at times 0 and 8 post IR respectively. ****, p<0.0001; ***, p<0.001 *, p<0.05; two-tailed Mann-Whitney test. Histograms show the percentage (mean±SD, N=5) of unrepaired γH2AX foci per cell after 8 hours of IR compared to time 0 of each condition in G2-phase. *, p<0,05; two-tailed Paired t-test. **(D)** Box and whiskers (5-95 percentile) plots show distribution of γH2AX foci per cell at G1-phase. Time 0 and 8 post IR. siRNA conditions as indicated. Data pooled from 5 experiments. 300 and 150 cells scored per condition and experiment at times 0 and 8 post IR respectively. ****, p<0.0001; ***, p<0.001 *, p<0.05; two-tailed Mann-Whitney test. Histograms show the percentage (mean±SD, N=5) of unrepaired γH2AX foci per cell after 8 hours of IR compared to time 0 of each condition in G1-phase. **(E)** Box and whiskers (5-95 percentile) plots show distribution of γH2AX foci per cell at S-phase. Time 0 and 8 post IR. siRNA conditions as indicated. Data pooled from 5 experiments. 150 cells scored per condition and experiment at times 0 and 8 post IR respectively. ****, p<0.0001; **, p<0.01 *, p<0.05; two-tailed Mann-Whitney test. Histograms show the percentage (mean±SD, N=5) of unrepaired γH2AX foci per cell after 8 hours of IR compared to time 0 of each condition in S-phase. *, p<0,05; two-tailed Paired t-test.

Interestingly, we noticed that the initial levels of γH2AX foci (time 0) were different in each condition, being significantly lower upon HDAC1 depletion at all cell cycle phases analysed (Figure 3C). This reduction was also observed in whole cell extracts by western blots, whereas no difference was detected in CHK1 S345 phosphorylation that, like γH2AX, is also driven by ATM upon IR (Figure S2A). Similarly, HDAC1 depletion caused a significant reduction in γH2AX but not in CHK1 S345 phosphorylation when treated with camptothecin (CPT), a topoisomerase I inhibitor that leads to replication-born DSBs (Figure S2B and C). These results indicate that the reduction in γH2AX foci was not specific for IR. Therefore, despite that CPT and IR induced similar levels of DNA damage in synchronized siHDAC1-treated and control cells, γH2AX signalling seemed reduced.

### HDAC1 depletion impairs γH2AX signalling after I-SceI induced DSBs

Our data showing a reduced level of γH2AX signalling after IR or CPT treatment suggests the possibility that a major defect of siHDAC1 cells in DDR is DSB signalling. To explore this further, we analysed γH2AX signalling by the more quantitative method of ChIP-qPCR using a chromosome-integrated construct where DSBs were induced enzymatically. For this, we constructed a U2OS-derived cell line (U2OS-Sce) in which we stably integrated a previously reported construct that consists in a GFP gene interrupted by the hygromycin resistance gene (Hyg) flanked by two I-SceI restriction sites and under the control of the CMV promoter (42). For DSB induction we generated a lentivirus carrying a I-SceI nuclease gene fused to the mCherry ORF and a glucocorticoid growth receptor (GR), to ensure retention of the I-SceI enzyme in the cytoplasm. DSBs were induced 24h after lentiviral transduction, by supplementing the transduced culture with dexamethasone (DEXA) to allow I-SceI-mCherry nuclear translocation. DEXA induced nuclear translocation of I-SceI-mCherry equally well in siC control and HDAC1-depleted cells, as determined by fluorescence microscopy (Figure 4A).

**Figure 4.**
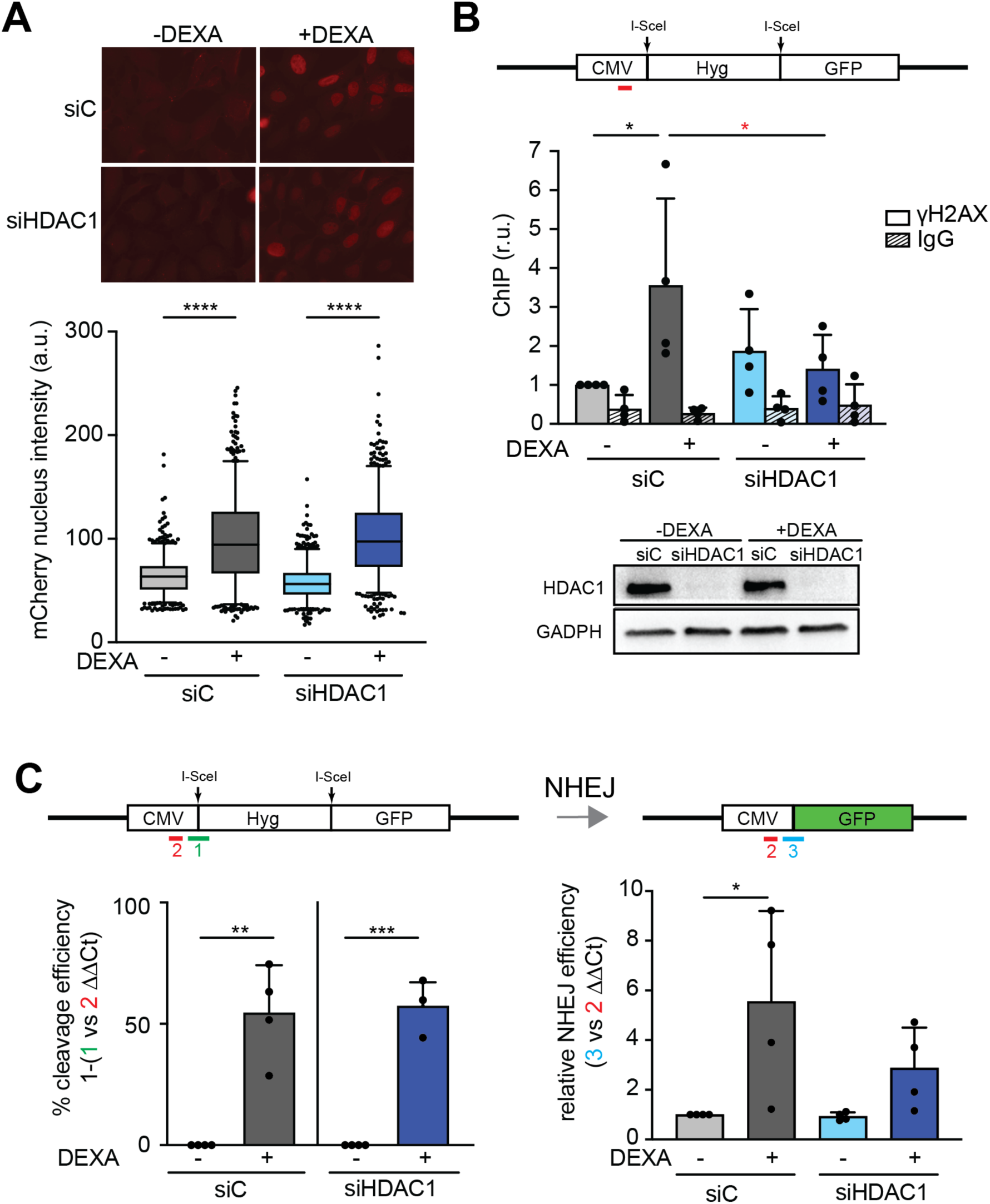
HDAC1 depletion impairs γH2AX signalling at I-SceI induced DSBs. **(A)** Images of mCherry (red) immunofluorescence of lentivirus transduced U2OS I-SceI cells. Box and whiskers (5-95 percentile) plots show distribution of mCherry nucleus intensity per cell. Data pooled from 4 experiments. >600 cells scored per condition. ****, p<0.0001; two-tailed Mann-Whitney test. Dexamethasone treatment and siRNAs as indicated. **(B)** ChIP analysis of γH2AX and IgG at 50bp away from the DSB site (red line). Histograms show the relative values of IP to siC -DEXA of each experiment (mean±SD, N=4). *, p<0.05; One-tailed Paired t-test. siRNAs and dexamethasone treatment as indicated. Representative Western-Blot of HDAC1 depletion in U2OS-ISce cell line. Anti HDAC1 and GADPH antibodies were used. siRNAs and DEXA treatment as indicated. **(C)** Histograms show the relative amount (mean±SD, N=4) of cleavage efficiency (green respect to red lines). ***, p<0.001; **, p<0.01; One-tailed Paired t-test. siRNAs and dexamethasone treatment as indicated. **(D)** Histograms show the relative amount (mean±SD, N=4) of NHEJ product (blue respect to red lines). *, p<0.05; One-tailed Paired t-test. siRNAs and dexamethasone treatment as indicated.

We next quantified γH2AX signalling by ChIP-qPCR at the I-SceI region after 6 hours of DEXA addition to the transduced cultures. In agreement with nuclear translocation of I-SceI-mCherry and DSB induction, a significant 4-fold increase in γH2AX signalling was detected in the siC control upon DEXA addition (Figure 4B). In contrast, no significant changes were observed in HDAC1-depleted cells. This result is consistent with defective signalling. To exclude the possibility of lower I-SceI cleavage efficiency upon HDAC1 depletion, we determined the cleavage efficiency of I-SceI by qPCR. For this, we first calculated the ratio of intact DNA (uncut DNA) by normalizing the qPCR signal obtained using specific primers that annealed at both sides of the upstream I-SceI restriction site versus the total qPCR signal obtained with primers annealing outside of the DSB region (Figure 4C). The cleavage efficiency was thus determined by subtracting the ratio of intact DNA from the total DNA. Approximately half of the molecules were cleaved in both siC control and siHDAC1 depleted cells indicating that the absence of HDAC1 did not affect I-SceI cleavage efficiency and confirming the role of HDAC1 in DSB signalling.

Since the two I-SceI*-*induced DSBs can lead to the loss of the hygromycin resistance gene when repaired by NHEJ, we also evaluated the NHEJ efficiency by qPCR with primers that amplify the resulting joined sequence. A significant 5.5-fold increase upon DEXA addition was detected in siC cells versus only a 3.4-fold in siHDAC1 cells, in agreement with the previously reported defect of HDAC1/2 depletion in NHEJ (23). This result fits with the conclusion that HDAC1 is required for DSB signalling, its depletion negatively impacting both HR, as determined by SCE, or NHEJ, as determined by this assay.

Our results therefore indicate that γH2AX signalling at I-SceI-induced DSBs is defective upon HDAC1 depletion, supporting a general defect in γH2AX signalling regardless of how DSBs are generated.

### Defective γH2AX signalling at LINE1-induced DSBs upon HDAC1 depletion

To further explore a putative role for HDAC1 in γH2AX signalling at DSBs, we designed a new system that allows to directly induce multiple DSBs in the genome. We used the U2OS SEC-C cell line expressing Cas9 (28) to target control (CT) or specific sgRNAs against the repetitive LINE1 (L1) sequences (43), which are present in 500,000 genomic copies (44). γH2AX and 53BP1 foci, which are known to require previous γH2AX foci formation (45), were measured by immunofluorescence 6 and 24 hours after L1-targeted sgRNA transfection. The levels of γH2AX and 53BP1 foci per cell significantly increased in both control and HDAC1-depleted cells upon L1-targeted transfection (Figure 5A and B). This confirms that multiple DSBs were caused by the expression of Cas9, thus validating our system.

**Figure 5.**
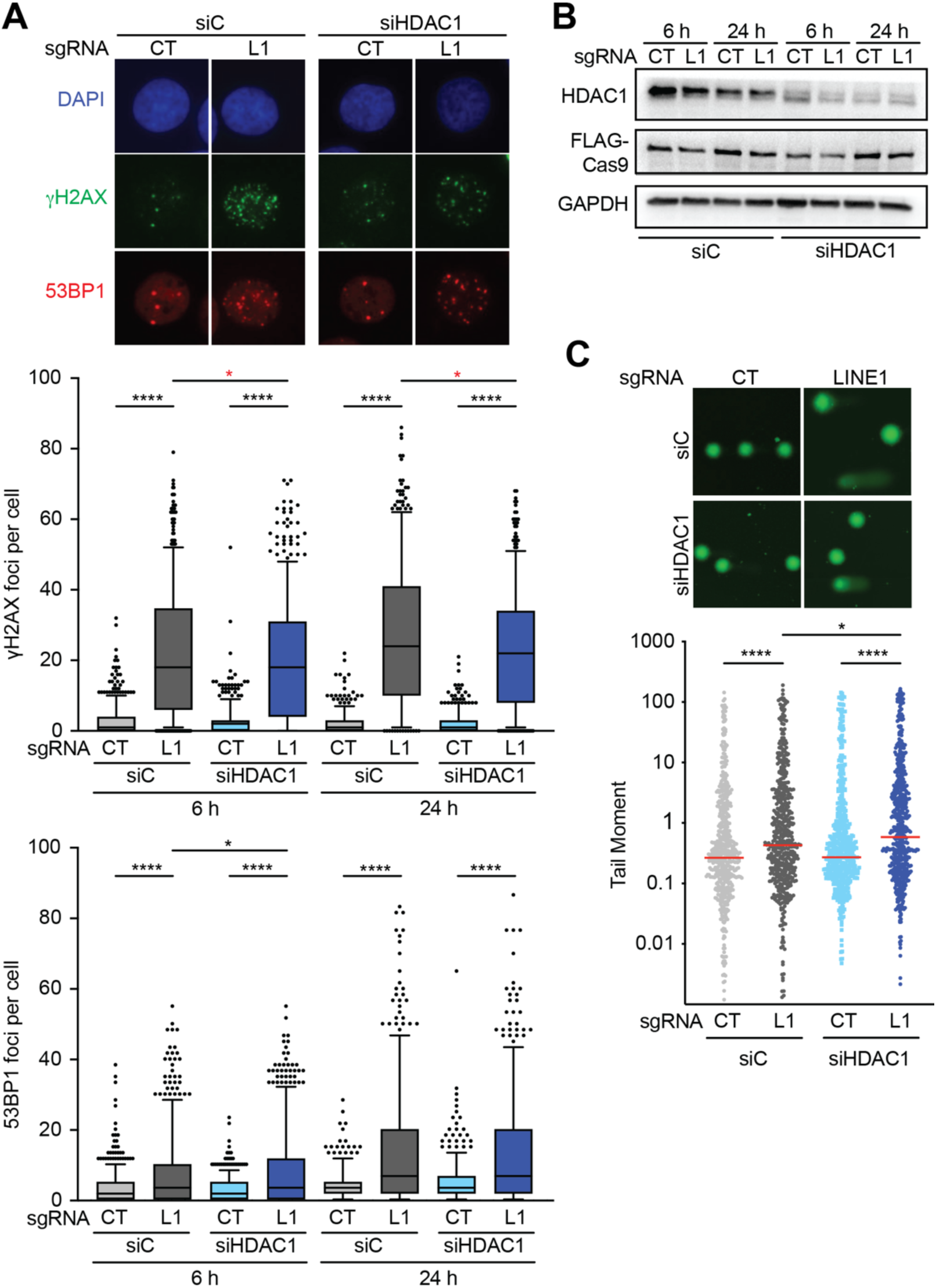
HDAC1 depletion impairs γH2AX signalling without affecting DSB formation at LINE1 sites. **(A)** Representative images of immunofluorescence with γH2AX (green), 53BP1 (red) antibodies and DAPI (blue) 6 hours after of L1 or CT sgRNA transfection. siRNAs and sgRNAs as indicated. Box and whiskers (5-95 percentile) plots show distribution of γH2AX and 53BP1 foci per cell respectively. siRNAs, sgRNAs and times post transfection as indicated. Data pooled from 4 experiments. 200 cells scored per condition and experiment. ****, p<0.0001; *, p<0.05; two-tailed Mann-Whitney test. **(B)** Representative Western-Blot of LINE1 (L1) and untargeted (CT) sgRNA transfection after Cas9 induction. Anti HDAC1, FLAG and GAPDH antibodies were used. sgRNAs and times post transfection as indicated. **(C)** Representative images of single-cell alkaline gel electrophoresis (comet assay). sgRNAs and times post transfection as indicated in both panels. Scatter dot plot shows tail moment distribution. Red line indicates the median. Data pooled from 3 experiments. 200 cells scored per condition and experiment. ****, p<0.0001; **, p<0.01; *, p<0.05; two-tailed Mann-Whitney test.

Notably, while the average number of 53BP1 foci per cell was similar or higher in HDAC1-depleted cells, the average number of γH2AX foci per cell was significantly lower (Figure 5A), supporting a defect in γH2AX signalling upon HDAC1 depletion. To confirm the massive cleavage of LINE1 elements in both control and HDAC1-depleted cells, we performed single-cell electrophoresis (comet assay) as a direct measurement of DNA breaks (Figure 5C). There was a strong increase in the comet tail moment in both cell lines upon L1-targeted transfection, with a higher increase in siHDAC1-treated cells. This excludes the possibility that reduction in γH2AX was due to lower levels of DSBs. In this case, the cleavage efficiency, estimated by qPCR as above, was low in both siC- and siHDAC1-treated cells, with a large proportion of genomic LINE1 copies remaining uncut (Figure S3A). Still, this was enough to cause a two-fold increase in γH2AX levels in siC cells after L1-targeted transfection (Figure S3B). HDAC1-depleted cells already showed increased spontaneous γH2AX levels (Figure S3B), consistent with its reported role preventing endogenous DNA damage (22,27,39,40,46). However, no further increase in γH2AX was detected after L1-targeted transfection. Altogether, these results indicate that DSBs can be induced equally throughout the genome by targeting Cas9 to endogenous LINE1 sequences in both control and HDAC1-depleted cells, but HDAC1 loss impairs the increase in γH2AX levels in the vicinity of such DSBs.

### HDAC1 loss impairs the DNA damage response after laser microirradiation

To directly assess the nuclear response to DNA damage, we induced DSBs by UV laser microirradiation and analysed the different DDR steps. By immunostaining with 53BP1 and γH2AX antibodies (Figure 6), we observed that the kinetics of 53BP1 recruitment was similar in control cells and in HDAC1-depleted cells, remaining at the breaks during the course of the experiment (60 min after the irradiation). In contrast, γH2AX intensity was significantly reduced ∼1.5 times in HDAC1-depleted cells as compared to control cells, already 15 min after laser microirradiation. The signal slowly declined in siC control cells because of the active repair, but it did not significantly change over time in HDAC1-depleted cells, consistent with a lower repair efficiency. These results indicate that HDAC1 depletion impairs γH2AX signalling also after laser microirradiation without affecting 53BP1 recruitment, which is known to happen independently of γH2AX (45).

**Figure 6.**
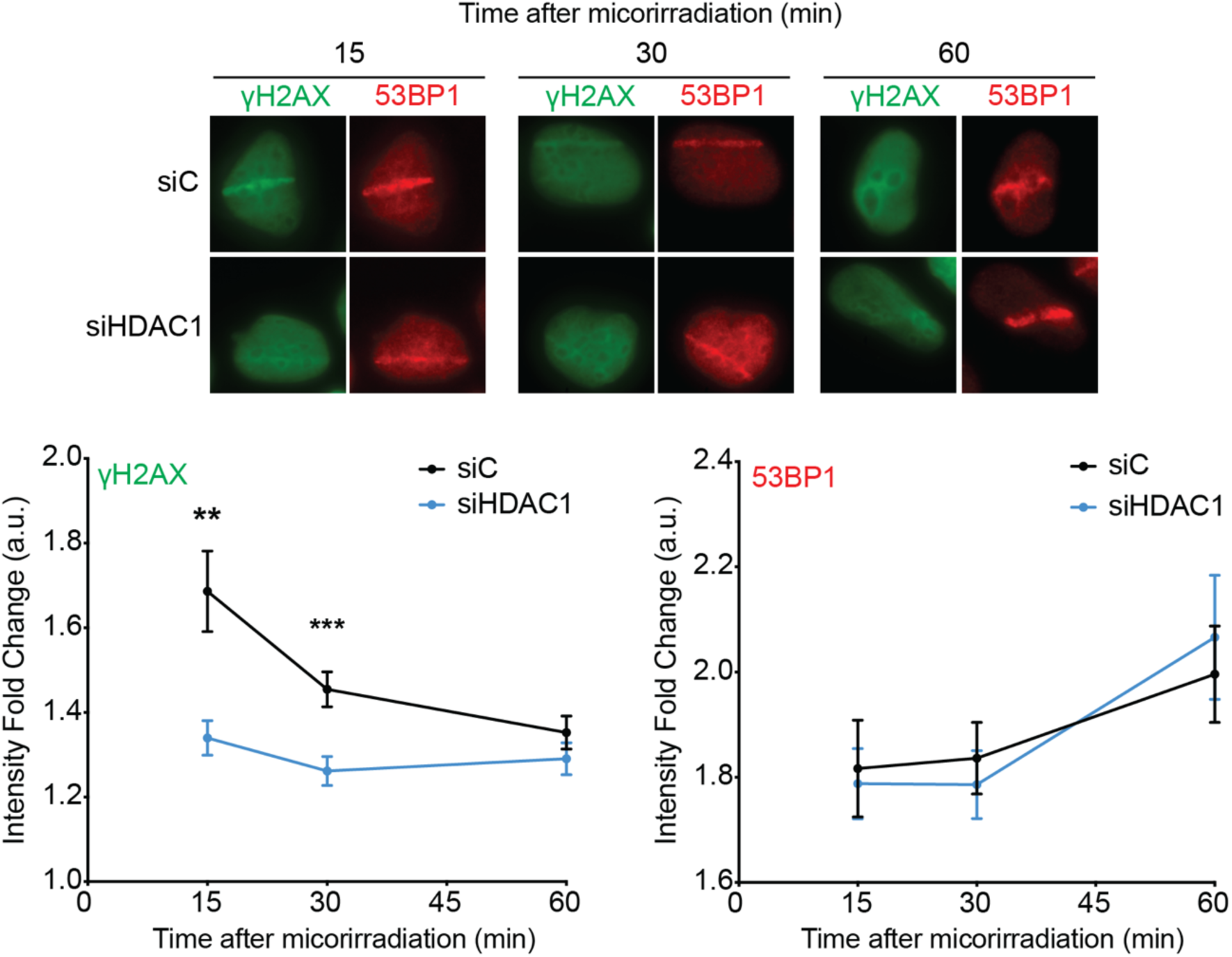
HDAC1 depletion impairs γH2AX signalling after laser microirradiation. Images of immunostaining with γH2AX (green) and 53BP1 (red) antibodies. siRNAs and times after laser microirradiation as indicated. Lower panel. Quantification of γH2AX and 53BP1 intensity fold change at laser stripes respectively. XY diagrams show mean±SEM (N=2, 20 cells per experiment) per condition and timepoint. ***, p<0.001; **, p<0.01; Two-way ANOVA test.

Next, we analysed the first stages of HR DSB repair by assessing the *in vivo* recruitment of the RPA70 subunit of the RPA factor, that coats the ssDNA generated after resection (5). HDAC1 depletion clearly reduced RPA70 recruitment levels at the laser stripe (Figure 7A), suggesting a defective or delayed DSB resection that is consistent with HR impairment. In contrast, a slight but consistent accumulation of the MRE11 subunit of the MRN complex was observed at the laser stripe in HDAC1-depleted cells (Figure 7B). This suggests that MRN, which promotes HR by removing the NHEJ factor KU70 from DSB ends (47), is retained at DSBs in the absence of HDAC1. Consistently, we observed similar levels of KU70 recruitment in HDAC1-depleted and siC control cells only at early time points (first 60 seconds), but HDAC1-depleted cells failed to reach the maximum levels of KU70 at the damaged DNA of the laser stripes that was observed in control cells (Figure 7C). These results agree with the defective NHEJ (Figure 4C) (23), suggesting a possible molecular basis for such a defect.

**Figure 7.**
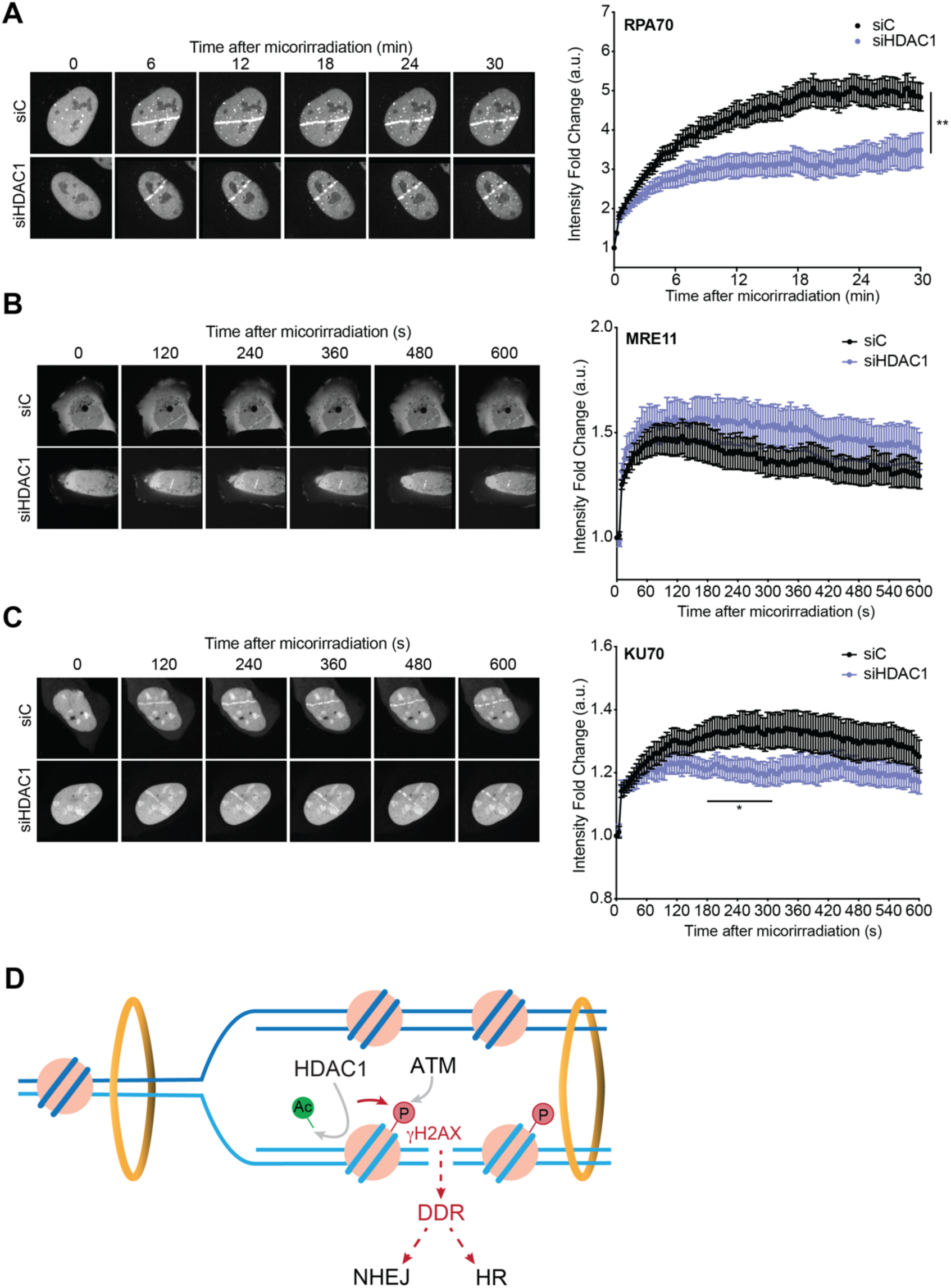
HDAC1 depletion affects the DNA damage response after laser microirradiation. **(A)** Images of U2OS RPA70-GFP cells upon laser microirradiation. siRNAs and times after laser microirradiation as indicated. Quantification of RPA70 intensity fold change at laser stripes. Intensity fold change at pre-laser time point (time 0) is normalized to 1 in both conditions. XY diagrams show mean±SEM (N=2, 10 cells per experiment) per condition and timepoint. **, p<0.01; Two-way ANOVA test. **(B)** Images of U2OS cells transfected with expression vector for MRE11-GFP protein upon laser microirradiation. siRNAs and times after laser microirradiation as indicated. Quantification of MRE11 intensity fold change at laser stripes. Intensity fold change at pre-laser time point (time 0) is normalized to 1 in both conditions. XY diagrams show mean±SEM (N=3, 10 cells per experiment) per condition and timepoint. **(C)** Images of U2OS cells transfected with expression vector for KU70-GFP protein upon laser microirradiation. siRNAs and times after laser microirradiation as indicated. Quantification of KU70 intensity fold change at laser stripes. Intensity fold change at pre-laser time point (time 0) is normalized to 1 in both conditions. XY diagrams show mean±SEM (N=2, 10 cells per experiment) per condition and timepoint. *, p<0.05; Two-way ANOVA test. **(D)** A model to explain the impact of HDAC1 driven deacetylation in the DNA damage response. HDAC1 mediates histone deacetylation around DSBs. This promotes ATM phosphorylation of H2AX. γH2AX triggers downstream DNA damage response (DDR) steps that end up in repair by NHEJ or HR, preferred when the sister chromatid is available such as behind replication forks. Upon HDAC1 inhibition or depletion, all downstream events are affected, from γH2AX to NHEJ and HR.

Altogether, our results show that HDAC1 loss impairs γH2AX signalling upon DNA damage leading to MRN retention and affecting DSB processing through both NHEJ and HR pathways.

## DISCUSSION

The analysis of the impact of different histone deacetylases in DSB repair reveals that HDAC1 activity is required for the efficient repair of DSBs by HR. Using genetics, molecular and cell biology approaches, we show that HDAC1 activity is necessary for γH2AX signalling, a first and key step in DSB repair, at DSBs induced by either IR genome-wide or by different nucleases at specific sites. Accordingly, HDAC1 depletion affects downstream events of the dynamic of repair factors at damaged sites (Figure 7D). As a consequence, cells lacking HDAC1 accumulate unrepaired DSBs and show hypersensitivity of cell survival to IR. HDAC1 and 2 were previously shown to impact NHEJ by their role on H3K56 acetylation (23). Here we focused on exploring the possible role of histone deacetylation in HR. We analysed the impact of inhibiting or depleting different HDACs on SCE, for which we first determined the impact of their depletion or inhibition in BLM-/- cells. As expected from the impaired dissolution of double Holliday junctions of cells lacking the BLM DNA helicase, the average number of SCEs detected in our experiments with BLM-/- cells (Figure 1) is in the same range as previous studies with other BLM-/- cells, such as CRISPR-generated BLM-/- RPE-1 cell lines (48), BLM-/- mouse embryonic fibroblasts (49), BLM-/- chicken DT40 cells (34) and cells from Bloom syndrome patients (32). Importantly, we detected a significant reduction SCEs after treatment with the TSA and Romidepsin HDAC inhibitors. Only HDAC1 depletion, but not depletion of HDAC2 and HDAC3 conferred the same SCE reduction levels in BLM-/- cells. This was confirmed also in BLM+/+ U2OS cells upon IR, indicating that HDAC1 depletion impairs HR regardless of the source of the damage.

An HR defect caused by HDAC inactivation was reported for budding yeast mutants in the HDAC1-related Rpd3L complex that also show SCE defects (19), suggesting that this is a conserved role of HDACs. As previously reported for yeast Rpd3L mutants (19), human HDAC1-depleted cells showed reduced levels of cohesins at chromatin (Figure S1). However, whereas yeast Rpd3L mutation confers a lower sister-chromatid cohesion that could explain the defective repair of replication-born DSBs, HDAC1 depletion did not induce a detectable premature sister-chromatid separation as it would be expected from a cohesion defect. This result agrees with previous reports in which premature sister chromatid separation was observed upon HDAC3 depletion, but not upon HDAC1 or HDAC2 depletion (50). Thus, rather than a defect in cohesion, our results support a crosstalk between two epigenetic marks, HDAC1-driven acetylation and phosphorylation of H2AX (Figure 7D), to explain the defective DSB repair. Consistently, and further in agreement with HDAC1 depletion impairing HR regardless of the source of the damage, we observed a defect in γH2AX signalling upon HDAC1 silencing in U2OS cells after IR (Figure 3 and S2), CPT (Figure S2) or UV laser microirradiation (Figure 6), as well by nucleases like I-SceI at specific sites (Figures 4) or Cas9 at LINE1 sequences along the genome (Figure 5). The fact that the effective cleavage of only a fraction of LINE1 endogenous targets (Figure S3) already allows to see the same effect as with the I-SceI or IR implies that the role of HDAC1 in γH2AX signalling is not sensitive to the nature of the sequences around the breaks, making it a general acting mechanism on DSBs. Thus, the mechanism by which HDAC1 affects SCR relies on its impact in γH2AX signalling. Indeed, γH2AX is known to promote HR to occur with the sister chromatid versus other templates (51).

γH2AX signal spreading around DSBs has been shown to depend on topologically associated domains, which are formed by cohesin loops through CBSs (52-54). It is therefore formally possible that the reduced levels of cohesin recruitment observed at CBSs upon HDAC1 depletion (Figure S1) could contribute to explain the HR defect by affecting γH2AX signalling. This is conserved in budding and fission yeast, in which HDAC mutants or inhibition by TSA were reported to reduce cohesion levels and induce sister chromatid separation (19,55). However, SCC1 depletion caused no defect in γH2AX signalling but rather an increase upon IR (Figure 3), as previously reported at nuclease-induced DSBs (56). Thus, we believe that the cohesion defect caused by HDAC1 depletion in human cells is minor and not enough to explain the HR-defective phenotype, which is mainly explained by the defective γH2AX signal.

Consistent with γH2AX signalling being a very early step in DDR, HDAC depletion affects downstream DSB repair events including the dynamic of RPA, MRN and KU70 at the damaged site (Figure 7). This agrees with the fact that γH2AX, although dispensable for the initial recruitment of DSB repair factors, is essential for the proper repair of DSBs by HR or NHEJ (45). Indeed, an NHEJ defect upon HDAC1 depletion was confirmed at I-SceI-induced DSBs (Figure 4), in agreement with the previously reported NHEJ defect (23). However, this had only been reported upon co-depletion of both HDAC1 and HDAC2 and attributed to H3K56 hyper-acetylation, which is only occurring when both HDAC1 and HDAC2 were depleted likely due to the facts that HDAC1 and HDAC2 share over 80% of the histone target sites and that the downregulation of one is compensated by the overexpression of the other one (23,57,58). In contrast, we observed no effect of HDAC2 depletion in SCE levels or in the percentage of unrepaired DSBs in any cell cycle phase (Figures 1 and 3). Thus, the effect described here is specific for HDAC1 function.

Supporting that the effect in HR of HDAC1 inactivation is direct, the HDAC inhibitor romidepsin had the same effect on SCEs (Figures 1 and 2). Moreover, HDAC1 is rapidly recruited to DSBs, earlier than proteins implicated in H2AX phosphorylation such as ATM or MDC1 (23,59,60). HDAC1 interacts with ATM (61). Thus, a priori its role in γH2AX signalling could be mediated by acting at ATM facilitating thus its recruitment and/or activation at DSBs. However, this possibility is ruled out because ATM acetylation drives its activation (62) and therefore, a putative ATM deacetylation driven by HDAC1 would have a negative effect in H2AX phosphorylation, rather than the positive effect observed here. Other HDAC1 non-histone targets that could contribute to its role in HR include RAD51. Indeed, HDAC1-mediated deacetylation of RAD51 has been reported to prevent its degradation (63). However, we favour a role for HDAC1 on histones. In this line, it is worth noticing that HDAC1 also interacts with the RSF1 chromatin remodeling in response to DSBs to induce the deacetylation of H2A(X)-K118 (64). This mark is important for both the ubiquitination of H2A-K119 that is associated with DNA damage-induced transcriptional repression (65), and for γH2AX signal spreading in G1 (64). Thus, it is possible that the effect that we see upon HDAC1 loss is due to defective H2AX deacetylation, that would precede H2AX phosphorylation.

Our results are of relevance in both proliferating and non-proliferating cells. γH2AX is a well-known tumour suppressor (45,66-68) and thus, HDAC1 mutations could also contribute to tumorigenesis and tumour development. Moreover, HDAC inhibitors are currently under extensive investigation for chemotherapy in certain cancer types and they are promising when combined with genotoxic agents (24,26). HDAC inhibitors not only cause DNA damage directly but also sensitize tumour cells to DNA-damaging agents and IR (25). HDAC inhibitors such as TSA and sodium butyrate (NaB) have been shown to radio sensitize human melanoma cell lines (69). Similarly, romidepsin has been shown to compromise the growth of osteosarcoma in vivo and in osteosarcoma cells such as the U2OS cells tested here (70). This increased sensitivity could be partially explained by the HR role uncovered here for HDAC1. HDAC1 inhibition could enhance genotoxic treatment of highly proliferating cancer cells, which repair DNA damage via HR. Along this line, the pan-HDAC inhibitor vorinostat has been shown to impair timely DNA damage repair, particularly in cancer cells (71). Despite the DNA damage repair defects caused by HDAC inhibition could be partially explained by the fact that it produces a transcriptional downregulation of DNA repair factors (27), our results support a direct contribution mediated by defective γH2AX signalling. Of note, impaired repair might also nourish the appearance of chemoresistance, as previously discussed (72). On the other hand, non-proliferating cells such as neurons mainly rely on NHEJ to counteract DNA damage that can lead to neurodegeneration in diseases such as Alzheimer and amyotrophic lateral sclerosis. In neurons, HDAC1 is known to interact at DSBs with the SIRT1 sirtuin, which is recruited in an ATM-dependent manner to promote the acetylation of HDAC1 itself (73), and FUS, which is phosphorylated by ATM (74) and recruited to DSBs to promote HDAC1 recruitment and DSB repair (59). Interestingly, and in agreement with our model, FUS depletion impaired not only HDAC1 recruitment but also γH2AX signalling (59). Thus, in the context of the brain, HDAC1 role in repair would be of relevance to prevent neurotoxicity. Consistently, dysregulation of HDAC1 activity has been reported to cause DSBs but also increased sensitivity to DNA damage (39). Our study, therefore, not only uncovers a specific role for HDAC1 in γH2AX signalling upon DNA damage, but also paves the way towards further investigations on HDAC1 as a therapeutic target.

## Supporting information

Supplemental filgures and tables

## Data availability

The data underlying this article will be shared on reasonable request to the corresponding author.

## ACKNOWLEDGEMENTS

We thank C. Lachaud, P. Huertas, J.A. Pintor-Toro and S. de Almeida for cell lines and plasmids.

## Author contributions

The study was conceived and supervised by A.A. and B.G.-G. J.- J.M.-G. performed the majority of the experiments, carried out the formal analyses, and validated the results with assistance from M.B.-F., S.S. and S. Silva. Methodology was developed by J.-J.M.-G. and S. Silva. Project administration and funding acquisition were undertaken by A.A. and B.G.-G. Visualization and figure preparation were conducted by J.- J.M.-G. and B.G.-G. The original draft of the manuscript was written by J.-J.M.-G., A.A., and B.G.-G., and all authors contributed to manuscript review and editing. All authors approved the final version of the manuscript.

