## Supplemental filgures and tables for "HDAC1 has a role in double strand break repair by regulating γH2AX signalling"

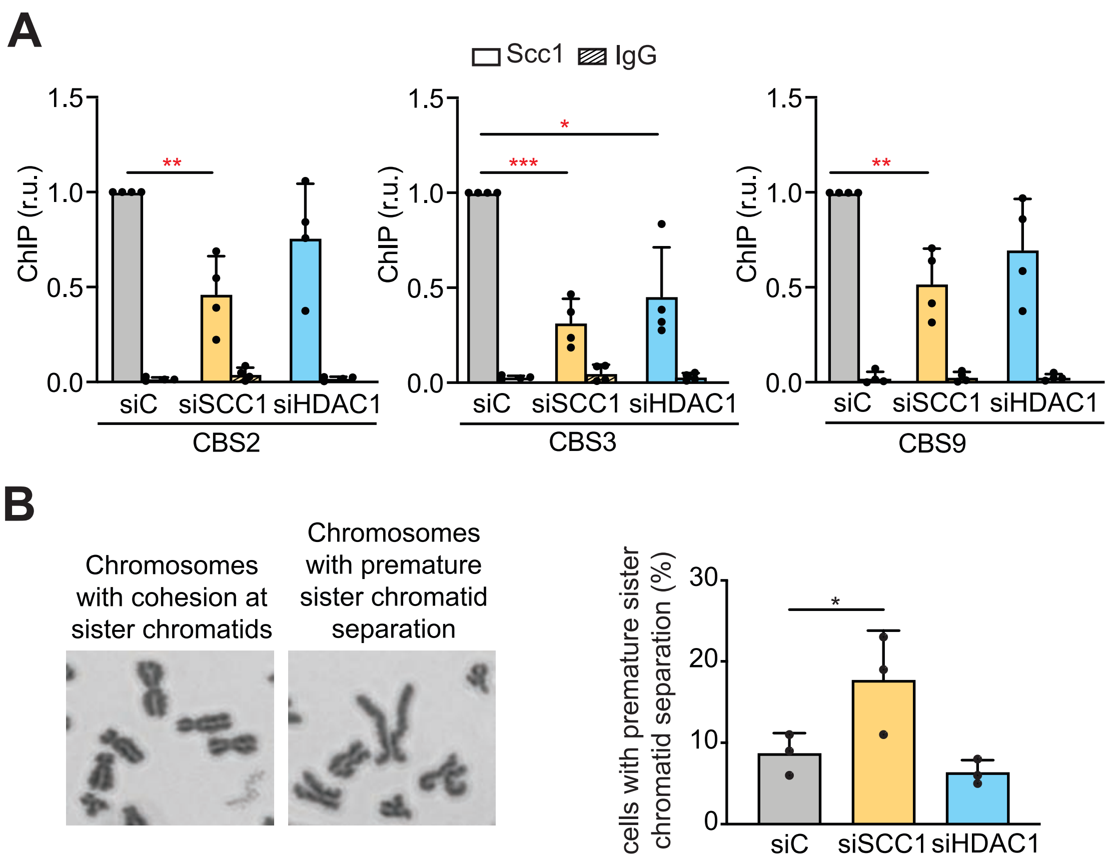


**Figure S1. HDAC1 depletion affects cohesion without inducing premature sister chromatid separation.**
**(A)** ChIP-qPCR analysis of SCC1 and IgG at Cohesin Binding Sites (CBS). Histograms show the relative values of IP to siC of each experiment (mean±SD, N=4). ***, $p<0.001$; *, $p<0.05$; two-tailed Paired t-test
**(B)** Images of normal and premature sister chromatid separation (PSCS). Histogram show the percentage (mean±SD, N=3) of PSCS cells. 100 metaphase per condition and experiment. siRNAs as indicated. *, $p<0.05$; two-tailed Paired t-test.

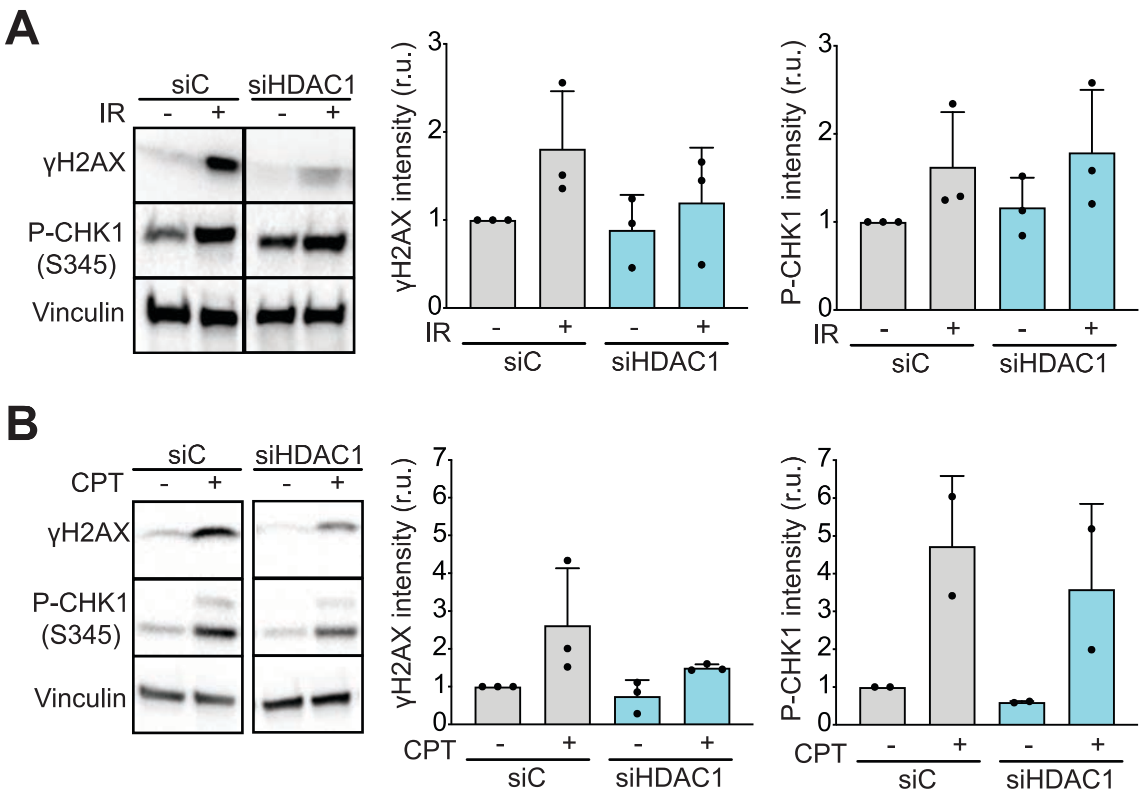


**Figure S2. HDAC1 depletion affects γH2AX signaling upon induced-DNA damage.**

**(A)** Representative Western-Blot of non-IR and IR (15 min post irradiation) U2-OS cells. Anti γH2AX, P-CHK1 (S345) and Vinculin antibodies were used. siRNA as indicated. Histograms show the quantification of γH2AX and P-CHK1 intensity (mean±SD, N=3) relative to NI siC of each experiment. *, $p<0.05$; two-tailed Paired t-test.

**(B)** Representative Western-Blot of CPT-treated U2-OS cells. Anti γH2AX, P-CHK1 (S345) and vinculin antibodies were used. siRNA as indicated. Histograms show the quantification of γH2AX and P-CHK1 intensity (mean±SD, N=3/2) relative to NT siC of each experiment.

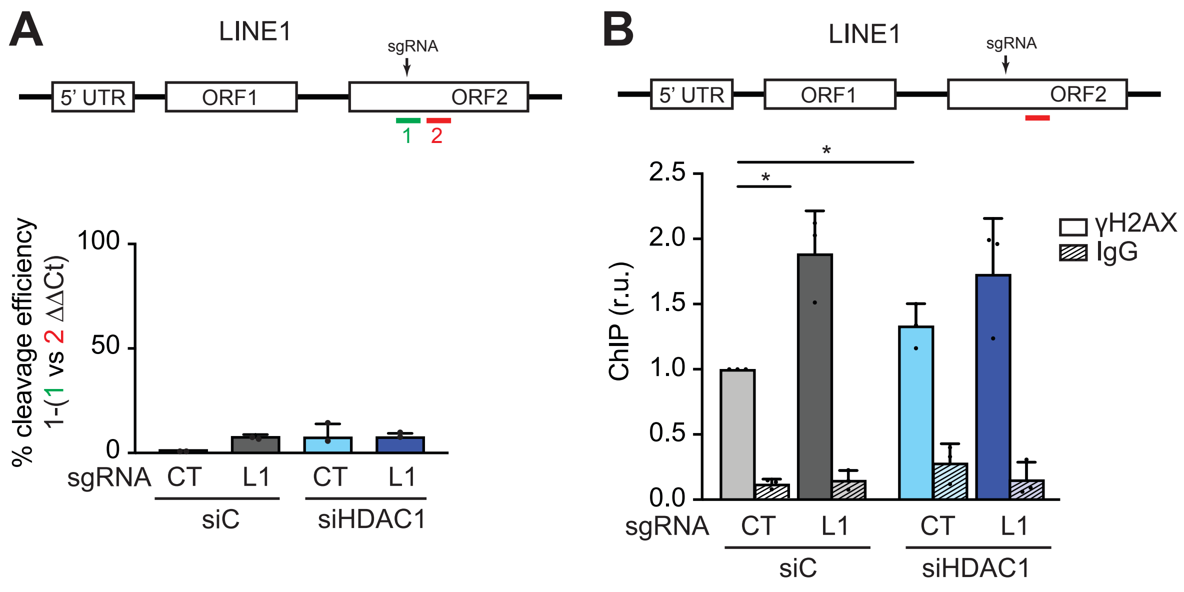


**Figure S3. HDAC1 depletion impairs γH2AX signaling at LINE1-induced DSBs.**

**(A)** Histograms show the amount of 1 relative to 2 (mean±SD, N=2) at L1 elements. siRNA and sgRNA as indicated.

**(B)** ChIP analysis of γH2AX at 175 bp away from the DSB site (red line). Histograms show the relative values of IP to siC CT of each experiment (mean±SD, N=3). *, $p<0.05$; one-tailed Paired t-test.

### Table S1- Reagents and Tools

| Reagent/resource | Reference or source | Identifier or cat number |
|---|---|---|
| **Antibodies** | | |
| Rabbit anti Histone H3 (acetyl K9 + K14 + K18 + K23 + K27) | Abcam | cat# ab47915; RRID:AB_873860 |
| Rabbit anti H3 | Merck | cat# H0164; RRID:AB_532248 |
| Mouse anti HDAC1 | Santa Cruz | cat# sc-81598; RRID:AB_2118083 |
| Mouse anti HDAC2 | Abcam | cat# ab12169; RRID:AB_2118547 |
| Mouse anti HDAC3 | Cell Signaling | cat# 3949; RRID:AB_2118371 |
| Mouse anti SCC1 | Merck | cat# 05-908; RRID:AB_11214315 |
| Rabbit anti SCC1 (ChIP) | Abcam | cat# ab217678; RRID:AB_2920658 |
| Mouse anti Vinculin | Merck | cat# V9264; RRID:AB_10603627 |
| Rabbit anti GAPDH | Cell Signaling | cat# 2118; RRID:AB_561053 |
| Mouse anti FLAG | Merck | cat# F3165; RRID:AB_259529 |
| Mouse anti Phospho-Histone H2A.X (Ser139) (WB) | Merck | cat# 05-636; RRID:AB_309864 |
| Mouse anti Phospho-Histone H2A.X (Ser139) (IF) | BioLegend | cat# 613402; RRID:AB_315795 |
| Rabbit anti Phospho-Histone H2A.X (Ser139) (ChIP) | Abcam | cat# ab2893; RRID:AB_303388 |
| Rabbit anti Phospho-CHK1 (S345) | Cell Signaling | cat# 2348; RRID:AB_331212 |
| Rabbit anti 53BP1 | Novus Biologicals | cat# NB100-304; RRID:AB_10003037 |

| | | |
|---|---|---|
| Rabbit anti IgG | Merck | cat# M7023; RRID:AB_260634 |
| Mouse anti IgG | Merck | cat# I8765; RRID:AB_1163672 |
| Goat anti-mouse IgG-HRP | Invitrogen | cat# A4416; RRID:AB_258167 |
| Goat anti-rabbit IgG-HRP | Invitrogen | cat# A0545; RRID:AB_257896 |
| Goat anti-mouse IgG (Alexa Fluor 488) | Invitrogen | cat# A11029; RRID:AB_2534088 |
| Goat anti-rabbit IgG (Alexa Fluor 568) | Invitrogen | cat# A11011; RRID:AB_143157 |
| **Chemicals, peptides, and recombinant proteins** | | |
| Protease inhibitor cocktail | Roche | cat# 11873580001 |
| Phenyl-methylsulfonyl fluoride (PMSF) | Fisher Scientific | cat# ICN800263 |
| TSA | Sigma | cat# T8552 |
| Romidepsin | Selleckchem | cat# FR228 |
| DAPI | Merk | cat# 32670 |
| Hoechst 33258 | AnaSpec | cat# 83219 |
| Dexamethasone | Sigma aldrich | cat# D4902 |
| Caffeine | Sigma aldrich | cat# C0750 |
| KaryoMAX colcemid | Invitrogen | cat# 15212012 |
| Proteinase K | Roche | cat# 3115836001 |
| Camptothecin | Sigma aldrich | cat# 390238 |
| **Critical commercial assays** | | |
| Click-iT plus EdU Alexa Fluor 647 Imaging Kit | Invitrogen | cat# C10640 |
| CometAssay™ kit | Trevigen | cat# 4250-050-K |
| iTaq universal SYBR Green supermix | Bio-rad | cat# 172-5124 |
| **Experimental models: Cell lines** | | |
| U2OS | ECACC | 92022711 |

| | | |
|---|---|---|
| BLM -/- fibroblast | Coriell Institute | GM08505 |
| U2OS SEC-C | C. Lachaud | (Munoz *et al.*, 2014) |
| U2OS-ISce | This study | N/A |
| U2OS RPA70-GFP | P. Huertas | N/A |
| **Recombinant DNA** | | |
| pKU70-GFP | (Britton *et al*, 2013) | |
| pMRE11-GFP | (Wijnhoven *et al*, 2015) | |
| pCMV-VSV-G | Addgene | cat# 8454 |
| pCMV-DR8.91 | Addgene | cat# 8455 |
| pSIN-DUAL-mCherry-ISceI-GR | This study | N/A |
| **Software and algorithms** | | |
| LAS AX | Leica | N/A |
| Thunder DMi8 | Leica | N/A |
| NIS Elements 4.0 | Nikon | N/A |
| MetaMorph v7.5.1.0 | Molecular Probes | N/A |
| Comet-score (v. 1.5) | TriTek Corp. | N/A |
| ImageJ 1.53a | NIH | N/A |
| ImageJ2 v2.9.0/1.53t | NIH | N/A |
| Primer express 3.0 | Applied Biosystems | N/A |
| Image Lab 6.0.1 | Bio-Rad | N/A |
| Prism v10.0.3 | GraphPad | N/A |
| **Other** | | |
| Lipofectamine 2000 | Invitrogen | cat# 11668019 |
| Lipofectamine 3000 | Invitrogen | cat# L3000015 |
| RNAi Max Lipofectamine | Invitrogen | cat# 10601435 |
| Polybrene | Sigma aldrich | cat# TR-1003 |
| Protein A Dynabeads | Fisher Scientific | cat# 10001D |
| Streptavidin magnetic beads | Millipore | cat# LSKMAGT02 |

| | | |
|---|---|---|
| Blocking reagent | Roche | cat# 11096176001 |
| SuperSignal West Pico Plus | Fisher Scientific | cat# 34580 |
| Buffer tablets pH 6,8 for Weisse Buffer | Sigma aldrich | cat# 1113740100 |
| ProLong Gold AntiFade reagent | Fisher Scientific | cat# P36930 |
| SYBR Gold Nucleic Acid Gel Stain | Invitrogen | cat# S11494 |
| Amersham Protran 0.2mM Nitrocelllulose Blotting membrane | Merk | cat# 10600001 |
| 4-20% gradient SDS-PAGE Criterion$^{TM}$ TGX$^{TM}$ Precast Gels | Bio-Rad | cat# 5671094 |
| Triton$^{TM}$ X-100 | Sigma aldrich | cat# X100 |

**Table S2- siRNAs and Oligonucleotides**

| OLIGO | SOURCE | IDENTIFIER/ SEQUENCE |
|---|---|---|
| ON-TARGETplus Non-targeting Pool | Dharmacon | cat# D-001810-10 |
| ON-TARGETplus SMART pool HDAC1 | Dharmacon | cat# L-011763-00 |
| ON-TARGETplus SMART pool HDAC2 | Dharmacon | cat# L-003495-00 |
| ON-TARGETplus SMART pool HDAC3 | Dharmacon | cat# L-003496-00 |
| Alt-R® CRISPR-Cas9 tracrRNA | Integrated DNA Technologies | cat# 1072533 |
| Alt-R® CRISPR-Cas9 Negative Control crRNA #1 | Integrated DNA Technologies | cat# 1072544 |
| Alt-R® CRISPR-Cas9 crRNA LINE-1 | (Chen *et al.*, 2017) | AUU CUA CCA GAG GUA CAA GG [rG][rU][rU][rU][rU][rA][rG][rA][rG][rC][rU][rA][rU]- |
| SCC1 siRNA.1 | This study | GCCCAUGUGUUCGAGUGUA[dT][dT] |
| SCC1 siRNA.2 | This study | UACACUCGAACACAUGGGC[dT][dT] |
| BamHI-mCherry-Fw | This study | GCTGCTGGATCCGCCACCATGGACAACACCGAGG |
| NotI-GR-Rv | This study | GCTGCTGCGGCCGCTTATCTAGATCCAGTGGAGCCAAATTT |
| L1 DSB Fw | This study | CCAGGACCAGATGGATTCACA |
| L1 DSB Rv | This study | GTTTCAGAAGGAATGGTACCAGTTC |
| ChIP-Uncut L1 Fw | This study | CAATAAAATACTGGCAAACCGAATC |
| ChIP-Uncut L1 Rv | This study | AGGGATGAAGCCCATTTGATC |
| I-SceI DSB Fw | | GCACCAAAATCAACGGGACT |
| I-SceI DSB Rv | (Chen *et al.*, 2017) | AAAGCACGAGATTCTTCGCC |
| I-SceI NHEJ Fw | (Chen *et al.*, 2017) | GCACCAAAATCAACGGGA |
| I-SceI NHEJ Rv | (Chen *et al.*, 2017) | GCTGAACTTGTGGCCGTTTA |
| ChIP-Uncut I-SceI Fw | (Chen *et al.*, 2017) | CCAAGTCTCCACCCCATTGA |
| ChIP-Uncut I-SceI Rv | (Chen *et al.*, 2017) | GCGGATCTGACGGTTCACTA |
| CBS2 Fw | (Panigrahi *et al.*, 2011) | AGTACACAGAGCGTAGCCTCC |
| CBS2 Rv | (Panigrahi *et al.*, 2011) | CCTCTAGTGGCTGATGCAGG |
| CBS3 Fw | (Panigrahi *et al.*, 2011) | CTCACTTCAGGTGGCAGAGG |
| CBS3 Rv | (Panigrahi *et al.*, 2011) | ATGGTCCAGTGTGGCATTC |
| CBS9 Fw | (Panigrahi *et al.*, 2011) | CAGCTCTGTGTCCTGTCTTATCC |
| CBS9 Rv | (Panigrahi *et al.*, 2011) | CAGCTATAATTGATGAAGAGGCG |
